# Research on the Mechanism of Lphn1 Knockout in Inhibiting Colorectal Cancer

**DOI:** 10.1101/2024.08.07.606975

**Authors:** Yi Wang

## Abstract

Decades ago, colorectal cancer was rarely diagnosed. Today, it is the fourth leading cause of cancer-related deaths worldwide, with nearly 90,000 fatalities each year. By analyzing single-cell data from tumor-bearing colorectal cancer model mice with Lphn1 knockout and wild-type Lphn1, we identified five key target genes for anticancer therapy: Ulbp1, Klrk1, Ccl6, Tlr4, Cd48, Prdm5, Vstm2a, Ret, Oas2, Hdac11 and Ptchd4, along with their corresponding cell types. Additionally, we discovered tumor-inhibiting cell subpopulations, including Cd244a_T_cells_subcluster_1, Cd48_Cd244a_NK_cells_subcluster_2, and C3_Macrophages_subcluster_1, which are potential candidates for therapeutic intervention. We propose that cancer-associated fibroblasts (CAFs) serve as the primary antigen presenters for MHC class I, providing antigens to macrophages, NK cells, and T cells to combat colorectal cancer. From a cellular perspective, the knockout of Lphn1 activates the anti-colorectal cancer functions of subpopulations of macrophages, NK cells, and T cells. Macrophages enhance antitumor immune activity by engaging the Ulbp1-Klrk1 receptor pair to activate NK cells. Additionally, macrophages activate downstream functions of T cells against colorectal cancer through CD48 signaling. After the knockout of Lphn1, macrophages are recruited by autocrine Ccl6 and Ccl6 secReted by CAFs. They exhibit high expression of Tlr4 and have the potential to transition into M1-type macrophages due to changes in their cellular state. After the knockout of Lphn1, the CAFs were reduced by half. CAFs, which are part of the cell network communication associated with tumor cells, typically play an immunosuppressive role. A reduction by half indicates that the immunosuppression in the Lphn1 knockout group has significantly decreased. This suggests that the efficacy of various cancer immunotherapy drugs can be significantly enhanced. In specific cell types, the four colorectal cancer-resistant transcription factors Irf7, Nr2f1 (with uncertain function), Bclaf1, and Irf2 co-localize with Tlr4, Cd48, Prdm5, and Oas2, respectively, and have been analyzed by pyscenic to interact and functionally contribute to the resistance against colorectal cancer. The four differential metabolic pathways between the Lphn1 group and the luc group are Arginine and proline metabolism, Histidine metabolism, Phenylalanine metabolism, and Riboflavin metabolism, with Arginine and proline metabolism being more active in the Lphn1 group and Histidine metabolism being more active in the luc group. CAF cells in tumors originate from CT26 cells. Additionally, Nr2f1 may serve as a potential therapeutic target, particularly as a transcription factor, for the treatment of colorectal cancer. These findings could open new avenues for the treatment of colorectal cancer and contribute to the development of personalized medicine.

## Introduction

Several decades ago, colorectal cancer was rarely diagnosed. Today, it is the fourth leading cause of cancer-related death in the world, with nearly 90,000 deaths annually. In addition to population aging and dietary habits in high-income countries, adverse risk factors such as obesity, lack of physical activity, and smoking have also increased the risk of developing colorectal cancer. Improved understanding of pathophysiology has expanded the treatment options for localized and advanced diseases, ultimately leading to personalized treatment plans. Treatment methods include endoscopic and surgical local resections, neoadjuvant radiotherapy and systemic therapy, surgical intervention for extensive local metastatic disease, localized ablative treatment for metastatic disease, palliative chemotherapy, targeted therapy, and immunotherapy. Although these new treatment approaches have doubled the overall survival rate for advanced disease to three years, the survival rate remains highest for patients without metastatic disease. Since this disease only presents symptoms in its advanced stages, organized screening programs are currently being implemented worldwide with the aim of improving early detection and reducing the incidence and mortality of colorectal cancer.

Colorectal cancer incidence varies across countries. Several factors are believed to contribute to this variation in incidence. In particular, among various socioeconomic status factors, low socioeconomic status is associated with an increased risk of colorectal cancer. In the United States, the incidence of colorectal cancer decreased from 60.5 per 100,000 individuals in 1976 to 46.4 per 100,000 in 2005. From 2003 to 2012, the incidence of colorectal cancer declined by approximately 3% each year. In 2017, there were 135,430 new cases of colorectal cancer in the U.S., with 50,260 deaths attributed to the disease. Although the overall incidence of colorectal cancer has declined, the incidence among individuals aged under 50 has increased by 2%. It is projected that by 2030, the incidence rates of colon and rectal cancer in patients aged 20 to 34 may rise by 90.0% and 124.2%, respectively. Approximately 35% of colorectal cancer cases in young individuals are believed to be related to hereditary colorectal cancer syndromes, though the reasons for the rising incidence remain unclear.

To investigate the complex causes of colorectal cancer, we performed single-cell sequencing analysis on tumor tissues from CT26 tumor-bearing mice. Single-cell sequencing allows for unprecedented resolution in exploring biological systems. The adhesion GPCR (Adgrl1/lphn1), known as Adgrl1 in humans and Lphn1 in mice, was previously shown to be highly expressed in human colorectal cancer as well as in most human tumors. However, after knocking out Lphn1 in mice, tumors in colorectal cancer model mice significantly shrank and lightened. This prompts our strong interest in understanding the mechanisms by which Lphn1 knockout inhibits colorectal cancer.

Therefore, we conducted single-cell sequencing analysis on tumor-bearing mice with and without Lphn1 knockout; the mice with Lphn1 knockout are referred to as the Lphn1 group, while the control group is referred to as the luc group. The single-cell sequencing analysis revealed significant differences in cell type abundance, and GO and KEGG enrichment analyses of differentially expressed genes indicated that macrophages regained their innate immune functions following Lphn1 knockout, NK cells exhibited activated immune functions and pro-tumor effects, while T cells in tumors from Lphn1-untouched mice undergone programmed cell death. Pseudotime analysis showed that macrophages underwent substantial functional changes post-Lphn1 knockout, acquiring the ability to positively regulate immune functions, modulate type 2 immune responses, and promote the proliferation of specific cell types, including T cells, mast cells, and epithelial cells(Supplementary Table 1). Cell communication analysis indicated that in Lphn1-knockout mice, a specific macrophage subpopulation (Subgroup 2) expressed Ulbp1, which activated natural killer (NK) cell-mediated cytotoxicity signaling pathways (both the ligand and receptor enriched within this pathway), enhancing the anti-tumor activity of NK cells(Supplementary Table 2). Anti-tumor active macrophages were recruited by self-secReted Ccl6 (Ccl6 expression was doubled compared to the luc group,Supplementary Table 3). The complement system of anti-tumor active macrophages contained fewer carcinogenic receptor-ligand signals compared to the luc group: C3 − (Itgam+Itgb2). Anti-tumor active macrophages secReted Tlr4, with secRetion levels 1.5-fold higher than the luc group, prompting the transformation of macrophages into M1-type anti-tumor macrophages(Supplementary Table 3). Simultaneously, Ccl6 secRetion by carcinoma-associated fibroblasts (CAF) within tumors of Lphn1-knockout mice was reduced by 0.5 times compared to the luc group, indicating CAFs’ role in recruiting macrophages with anti-tumor capabilities(Supplementary Table 4). Pseudotime analysis following Lphn1 knockout showed that NK cells underwent dramatic functional changes. Cell communication analysis indicated that NK cells secReted Cd48, a signal crucial for countering colorectal cancer. Pseudotime analysis of T cells suggested that the functional differences between cells with or without Lphn1 knockout were substantial. Cell communication analysis revealed that only in Lphn1-knockout cells did T cells express Cd48, indicating anti-tumor activity. Combined cell communication analysis of macrophages and T cells highlighted that only post-Lphn1 knockout, macrophages’ secReted Cd48 anti-tumor signal was received by T cells, executing the corresponding functional activation. Additionally, combined cell communication analyses between CAFs and macrophages, NK cells, and T cells indicated that regardless of Lphn1 knockout status, CAFs were able to present MHC1 anti-tumor signals to these three cell types, underscoring CAFs’ role as critical antigen-presenting cells in anti-tumor activity.

## Results

### 1.1 Results of Differential Signaling Analysis of Cell Types After Lphn1 Knockout through Single-Cell Analysis

Our single-cell sequencing was conducted using the Fudan Single-Cell Sequencing Platform, resulting in the analysis of 5,102 cells in the Lphn1 group (Lphn1 knockout tumor-bearing group) and 5,207 cells in the luc group (control group). The merged annotation results are shown in Figure 1a, categorizing the cells into nine different types: Monocytes, CAFs, NK cells, T cells, Granulocytes, Macrophages, Plasma cells, Mast cells, and B cells. The annotation results for the Lphn1 and luc groups are illustrated in Figure 1b. The frequency statistics of the cell types are depicted in Figures 1c and 1d, revealing significant differences in the quantities of five cell types: CAFs (the Lphn1 group had half the number compared to the luc group); NK cells (the Lphn1 group had three times more than the luc group); T cells (the Lphn1 group had 1.5 times more than the luc group); Granulocytes (the Lphn1 group had four times fewer than the luc group); and Macrophages (the Lphn1 group had half the number of the luc group). Such substantial differences in cell quantities indicate significant functional variations among these five cell types between the Lphn1 and luc groups.Figures 1e and 1f demonstrate a significant reduction in tumor size in mice after knockout of the Lphn1 gene.

**Figure 1.**
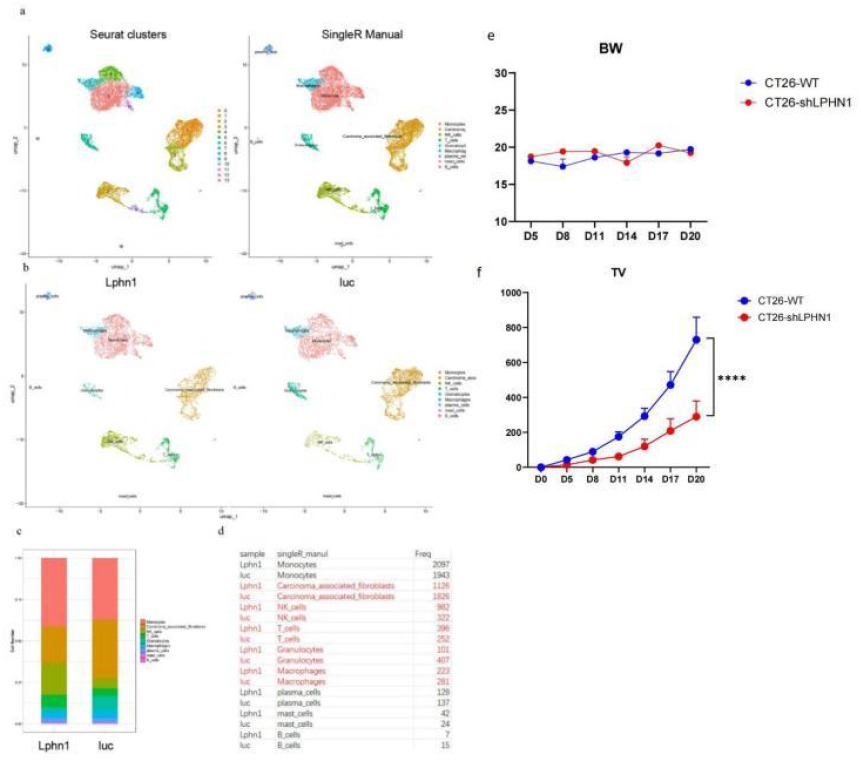
single-cell annotation results and cell count statistics for the lphn1 and luc groups. a. merged single-cell clustering map; b. annotation results for the lphn1 group and the luc group; c. proportional representation of cell counts in the lphn1 and luc groups; d. statistical table of cell count proportions in the lphn1 and luc groups. e. curve showing changes in body weight over time between gene knockout mice and control mice; f. curve showing changes in tumor size over time between gene knockout mice and control mice.

GO enrichment analysis of differentially expressed genes for each cell type revealed that Macrophages in the Lphn1 group restored innate immune functions, as shown in Figure 2a, with activation of biological processes such as innate immune response, immune response, immune system process, and regulation of immune system process. In contrast, the immune system of the luc group was not activated, as illustrated in Figure 2b.

**Figure 2.**
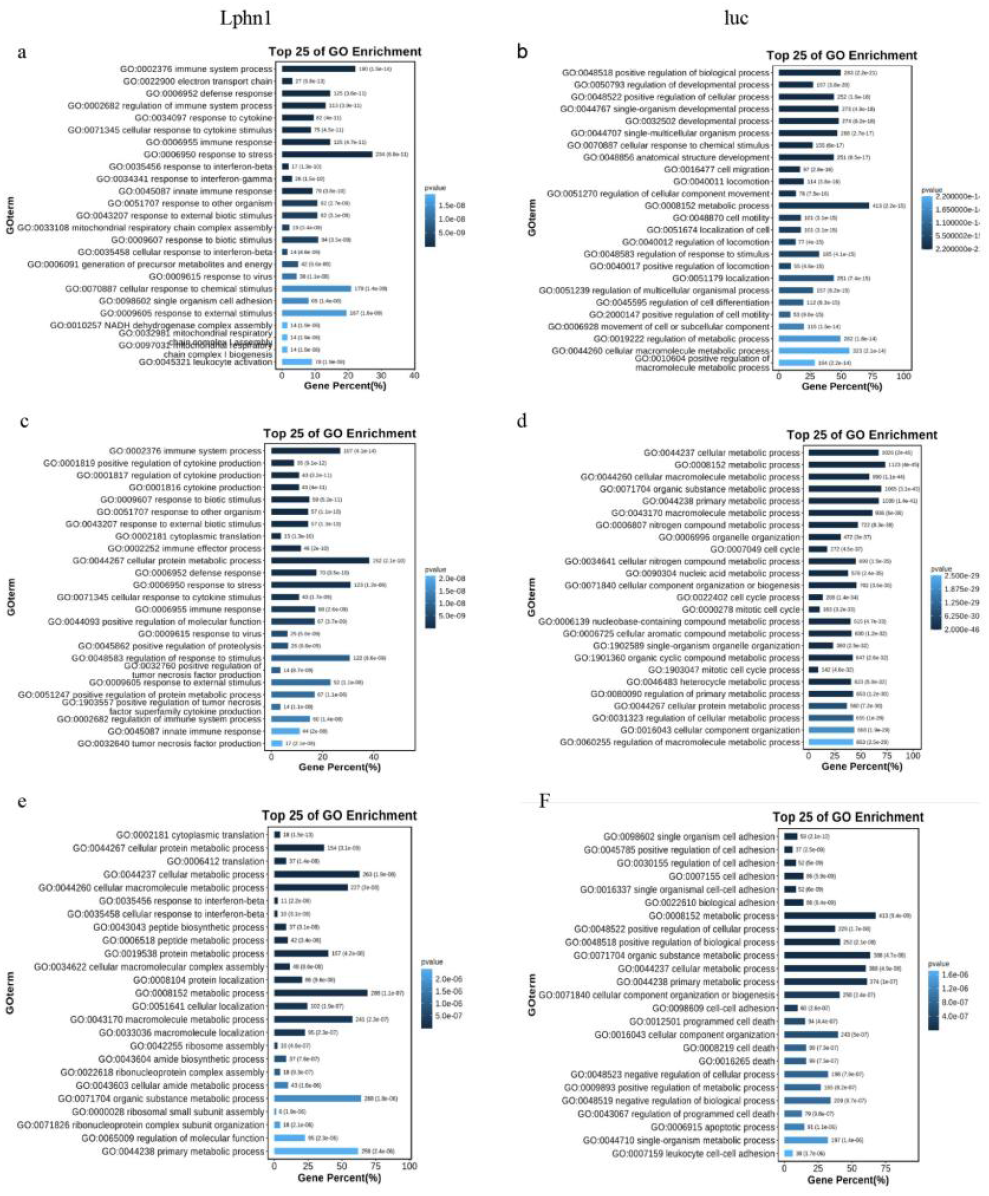
comparative analysis of go enrichment results for macrophages, nk cells, and t cells in the lphn1 and luc groups. a. biological process enrichment results for macrophages in the lphn1 group; b. biological process enrichment results for macrophages in the luc group; c. biological process enrichment results for nk cells in the lphn1 group; d. biological process enrichment results for nk cells in the luc group; e. biological process enrichment results for t cells in the lphn1 group; f. biological process enrichment results for t cells in the luc group.

NK cells in the Lphn1 group exhibited activated immune functions and enhanced anti-tumor capabilities, as shown in Figure 2c. Following the knockout of Lphn1, the biological processes activated in NK cells included immune system process, immune effector process, immune response, positive regulation of tumor necrosis factor production, positive regulation of tumor necrosis factor superfamily cytokine production, regulation of immune system process, innate immune response, and tumor necrosis factor production. In the luc group, however, NK cells did not exhibit these functions, as shown in Figure 2d.

The regulation of programmed cell death biological process was activated in T cells from the luc group, which may explain the lower T cell count compared to the Lphn1 group, as indicated in Figure 2e. Conversely, T cells in the Lphn1 group did not activate this biological process, as illustrated in Figure 2f.

### 1.2 Differences in Macrophages Between the Lphn1 and luc Groups

The pseudotime analysis results for the Lphn1 group indicated that cell states exhibited more branching, as shown in Figure 3a. In contrast, the tumor-bearing mice in the luc group displayed fewer branches in their Macrophages, suggesting that Macrophages in the Lphn1 group have functions that differ from those in the luc group. The cell state density map demonstrated that Macrophages in the Lphn1 group transitioned from one state to six states along the pseudotime, as illustrated in Figure 3c, while Macrophages in the luc group transitioned from one state to only two states, as shown in Figure 3d. This indicates a significant change in Macrophage functionality following the knockout of Lphn1.

**Figure 3.**
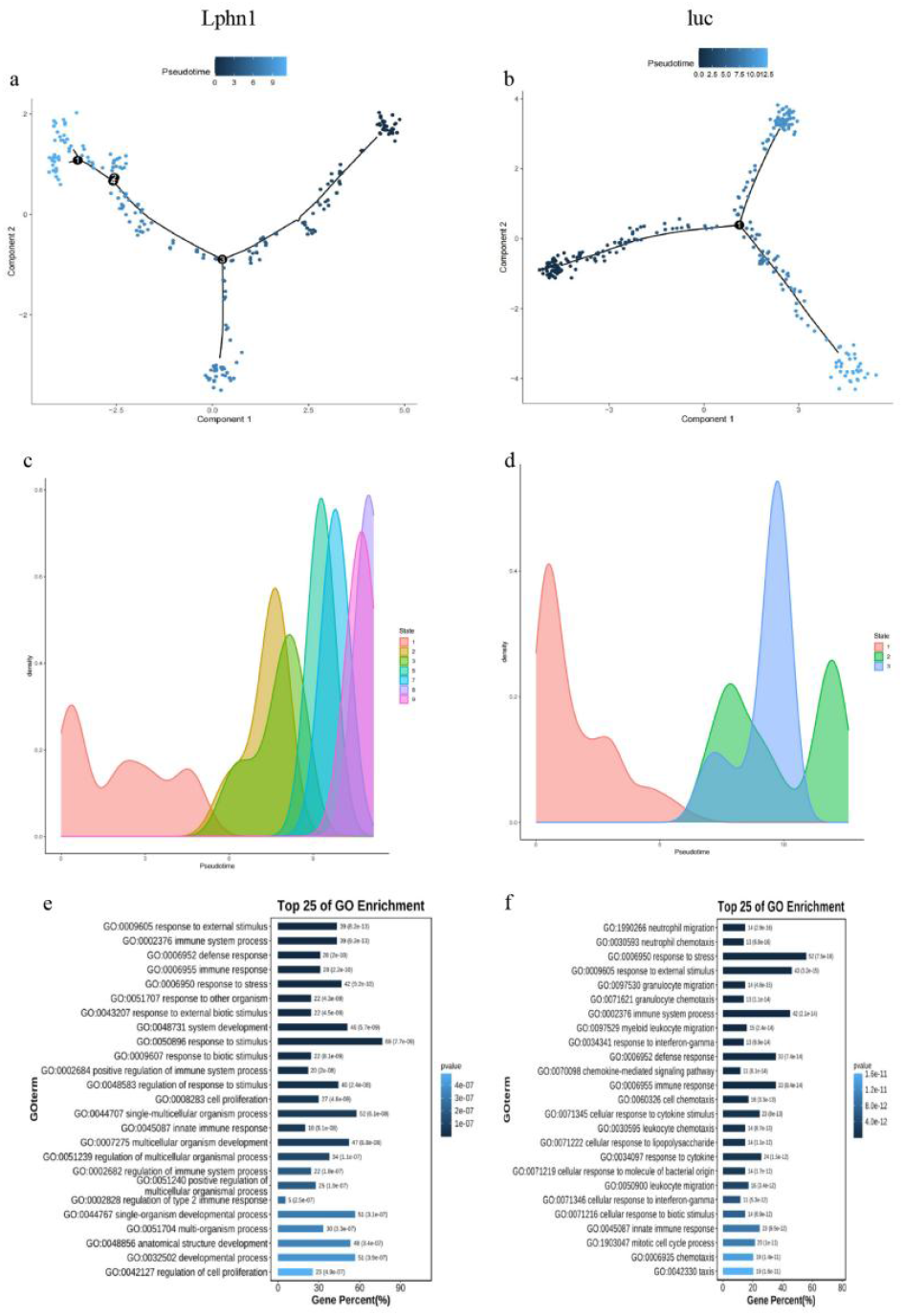
Pseudotime Analysis of Macrophages in the Lphn1 and luc Groups. a. Pseudotime branching trajectory of Macrophages in the Lphn1 group; b. Pseudotime branching trajectory of Macrophages in the luc group; c. Cell state density map of Macrophages in the Lphn1 group; d. Cell state density map of Macrophages in the luc group; e. GO biological pathway enrichment analysis of differentially expressed genes in Macrophages of the Lphn1 group; f. GO biological pathway enrichment analysis of differentially expressed genes in Macrophages of the luc group.

To further investigate, we conducted GO and KEGG enrichment analyses on the differentially expressed genes of Macrophages in the Lphn1 and luc groups. The results of the GO enrichment analysis revealed that Macrophages in the Lphn1 group possess the ability to positively regulate immune functions and modulate type 2 immune functions, as shown in Figure 3e (Lphn1 group) and Figure 3f (luc group). Additionally, they showed the capacity to promote the proliferation of specific cell types (Supplementary Table 1). The specific cell types include T cells, mast cells, and epithelial cells. Considering our finding that Tlr4 was significantly upregulated in the Lphn1 group compared to the luc group (1.5-fold increase, Supplementary Table 3), we have reason to speculate that Macrophages may polarize towards M1-type anti-tumor macrophages following the knockout of Lphn1.

Cell-to-cell communication analysis of Macrophage subpopulations in the Lphn1 and luc groups revealed that subpopulation 2 of Macrophages in the Lphn1 group represents the Ulbp1 subpopulation (referred to as Ulbp1_Macrophages_subcluster_2). This subpopulation activates the Natural Killer (NK) cell-mediated cytotoxicity signaling pathway through the Ulbp1-Klrk1 signaling, enhancing the anti-tumor activity of NK cells (Supplementary Table 2). This is illustrated in Figures 4a, 4b, 4c, and 4d.

**Figure 4.**
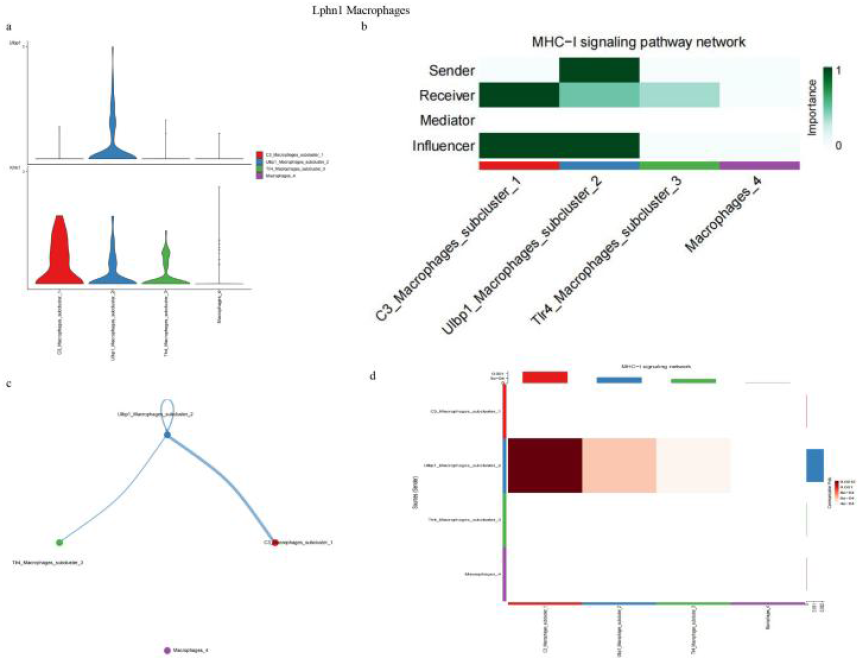
MHC1 Signaling Communication in Macrophages of the Lphn1 Group. a. Gene expression levels of MHC1; b. MHC1 signaling communication diagram; c. Intercellular communication structure of MHC1; d. Intercellular communication heatmap of MHC1.

Cell-to-cell communication analysis of Macrophage subpopulations revealed significant differences in the secRetion of Ccl (chemokine) between the Lphn1 and luc groups, as shown in Figures 5a, 5b, 5c, and 5d. This difference is primarily characterized by the self-secRetion of Ccl6 by Macrophages in the Lphn1 group, which recruits tumoricidal macrophages (presenting antigens to NK cells via Ulbp1-Klrk1 signaling to activate the Natural killer cell-mediated cytotoxicity pathway). The secRetion of Ccl6 by Macrophages in the Lphn1 group is twice that of the luc group (Supplementary Table 3). CCL6 is a chemokine that primarily acts as an attractant for macrophages, but it can also attract B cells, CD4+ lymphocytes, and eosinophils. Additionally, Ccl6 secRetion from Cancer-Associated Fibroblasts (CAFs) is 0.5 times higher in the Lphn1 group compared to the luc group (Supplementary Table 4), further contributing to the recruitment of anti-tumor macrophages.

**Figure 5.**
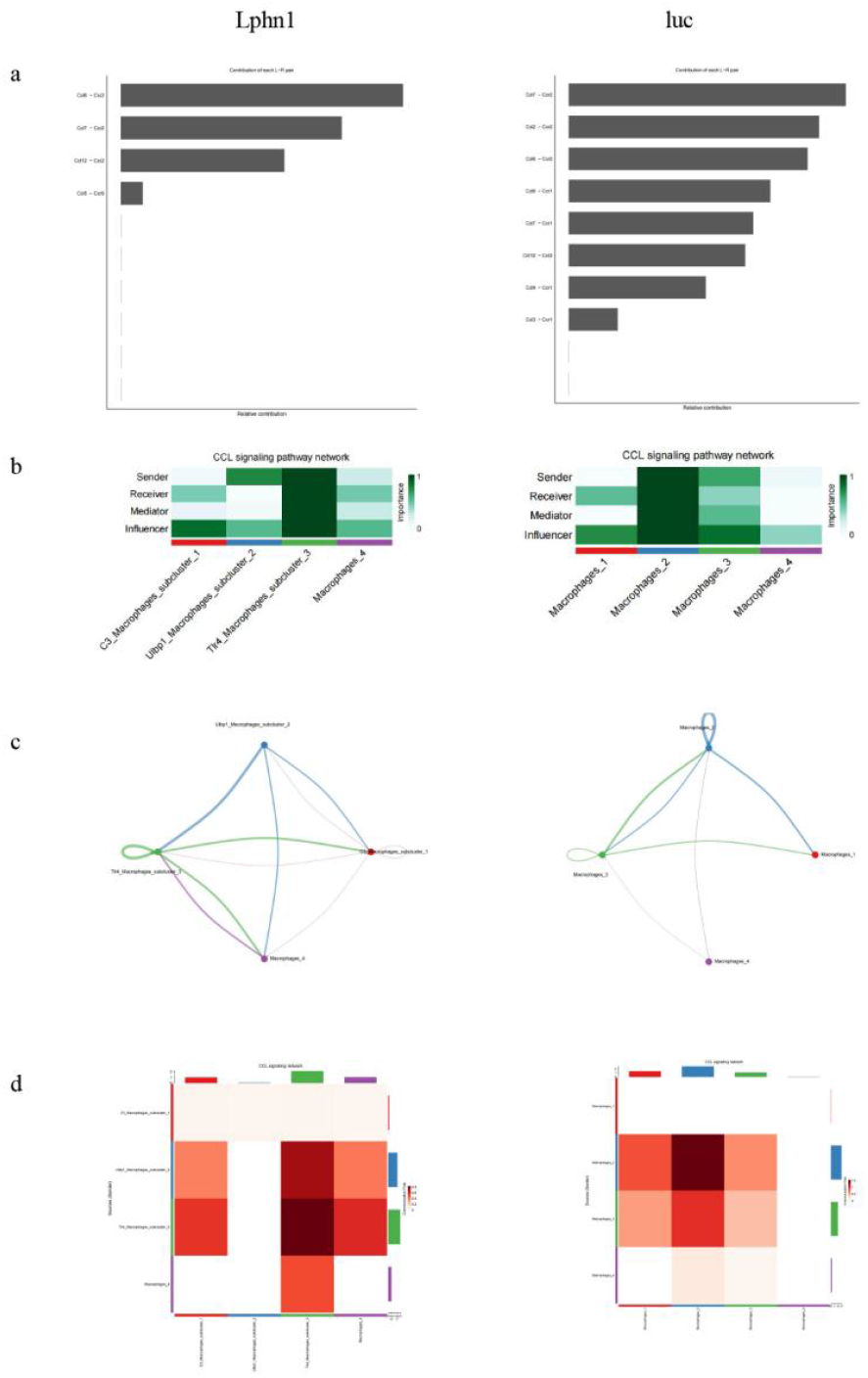
Comparison of Ccl Signaling Pathway Cell Communication between Macrophages in the Lphn1 and luc Groups. a. Comparison of Ccl signaling contribution differences between the two groups; b. Comparison of Ccl signaling communication differences between the two groups; c. Comparison of structural differences in Ccl signaling communication between the two groups; d. Heatmap comparison of differences in Ccl signaling communication between the two groups.

Meanwhile, in the complement system of Macrophages, the Lphn1 group shows a reduction in a cancer-associated receptor signal: C3 − (Itgam+Itgb2). We designate subpopulation 1 of Macrophages in the Lphn1 group as C3_Macrophages_subcluster_1, as illustrated in Figures 6a, 6b, 6c, and 6d.

**Figure 6.**
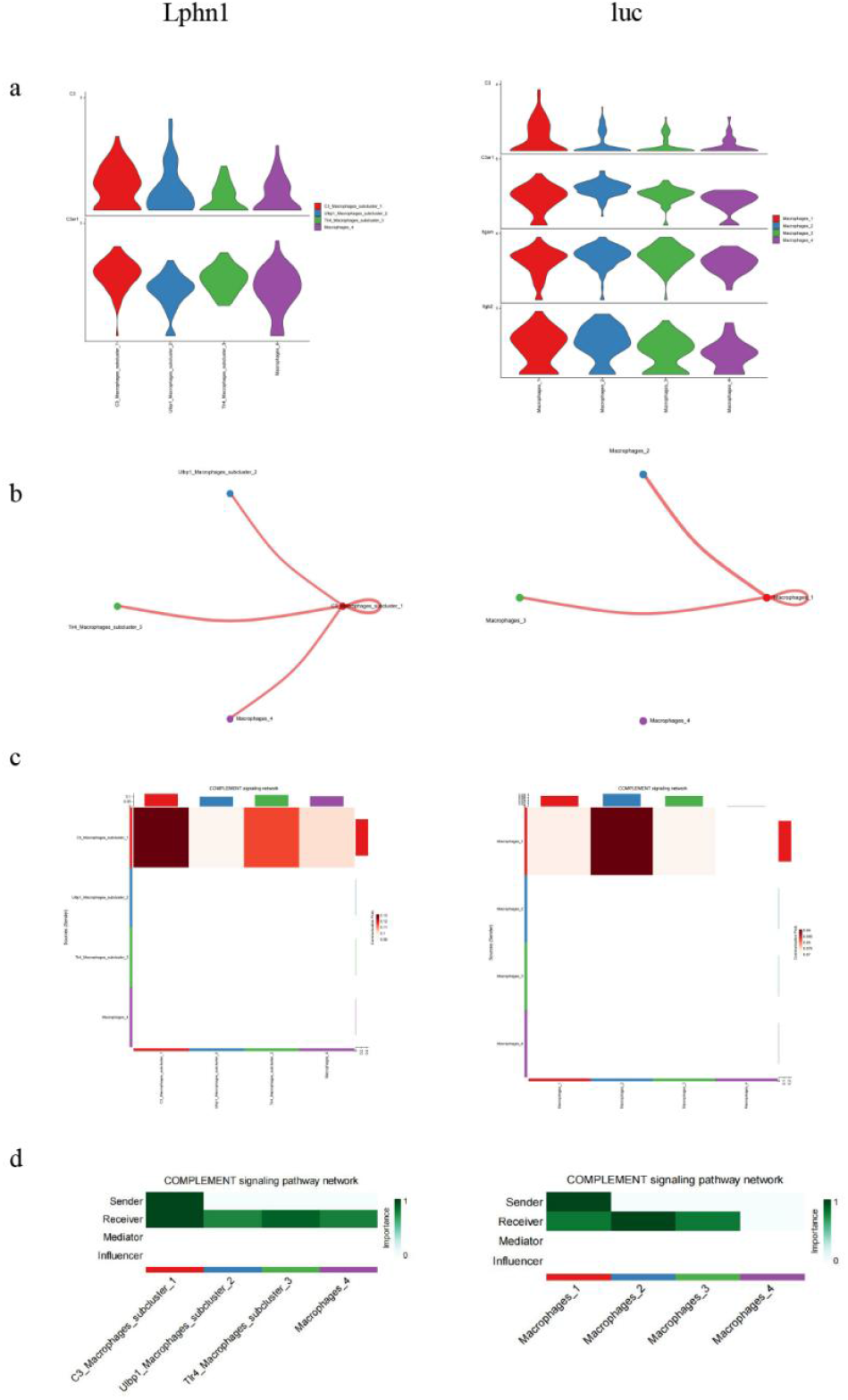
Differences in Complement Signaling Communication between the Lphn1 and luc Groups. a. Comparison of gene expression differences in complement signaling between the two groups; b. Comparison of structural differences in complement cell signaling communication between the two groups; c. Heatmap comparison of differences in complement signaling communication between the two groups; d. Comparison of signaling differences in complement cell communication between the two groups.

Tlr4 is significantly upregulated in the Lphn1 group compared to the luc group (increased by 0.5 times, Supplementary Table 3), which promotes the polarization of Macrophages in the Lphn1 group towards M1-type anti-tumor macrophages. We designate subpopulation 3 of Macrophages in the Lphn1 group as Tlr4_Macrophages_subcluster_3.

### 1.3 Differences in NK Cells between the Lphn1 and luc Groups

Pseudotime analysis of NK cells from the Lphn1 and luc groups revealed significant differences in their pseudotime cell states, as shown in Figures 7a and 7b. Notably, the cell density plot demonstrates a substantial difference, indicating that there are significant functional disparities in NK cells between the Lphn1 and luc groups.

**Figure 7.**
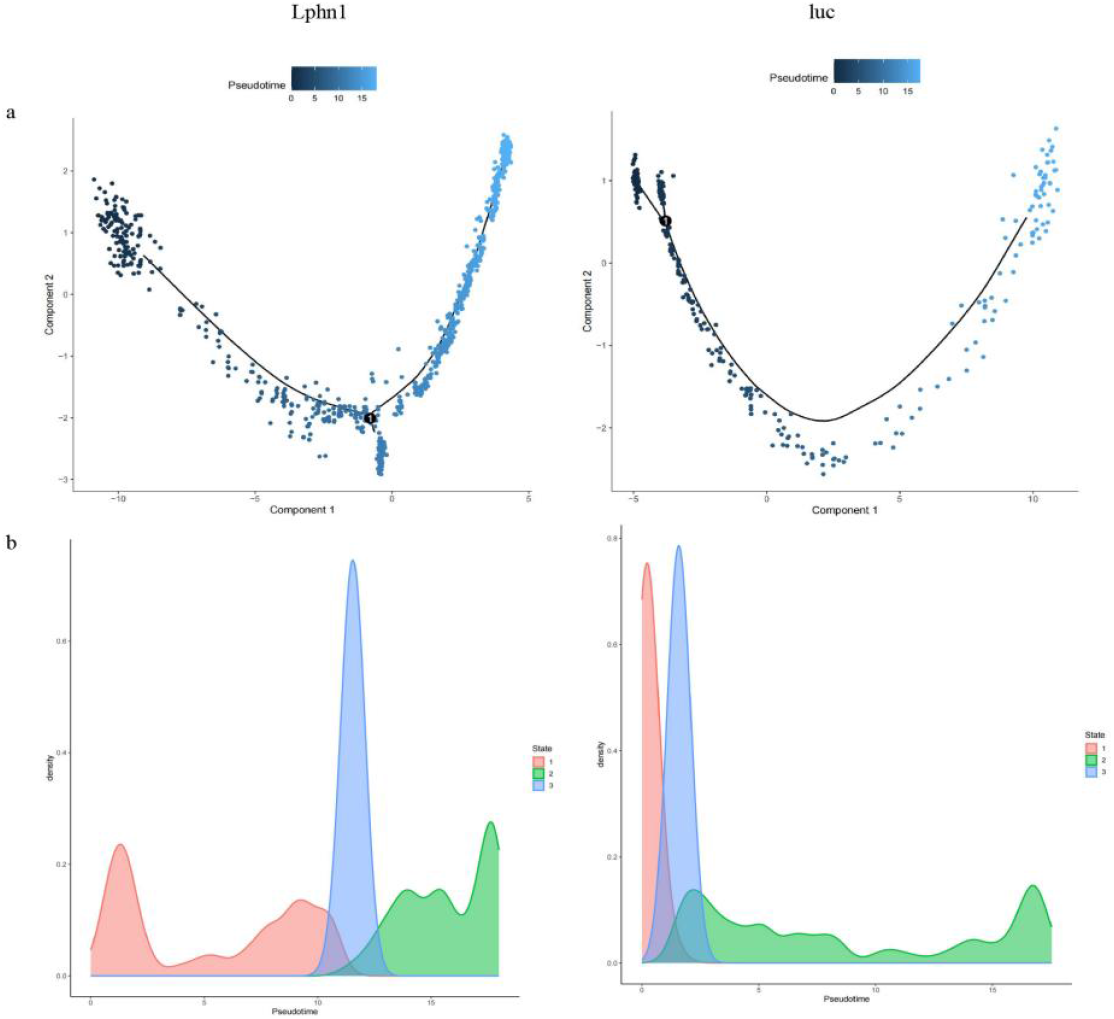
Pseudotime Analysis of NK Cells in the Lphn1 and luc Groups. a. Pseudotime branching trajectory of NK cells in the two groups; b. Cell state density plot of NK cells in the two groups.

The cell communication analysis of NK cells in the Lphn1 and luc groups revealed that Cd48 signaling was expressed only in the Lphn1 group, which is associated with anti-colorectal cancer effects, as shown in Figures 8a, 8b, 8c, and 8d. We designate subpopulation 2 of NK cells in the Lphn1 group as Cd48_Cd244a_NK_cells_subcluster_2, and subpopulations 4 and 5 as Cd48_NK_cells_subcluster_4 and Cd48_NK_cells_subcluster_5, respectively.

**Figure 8.**
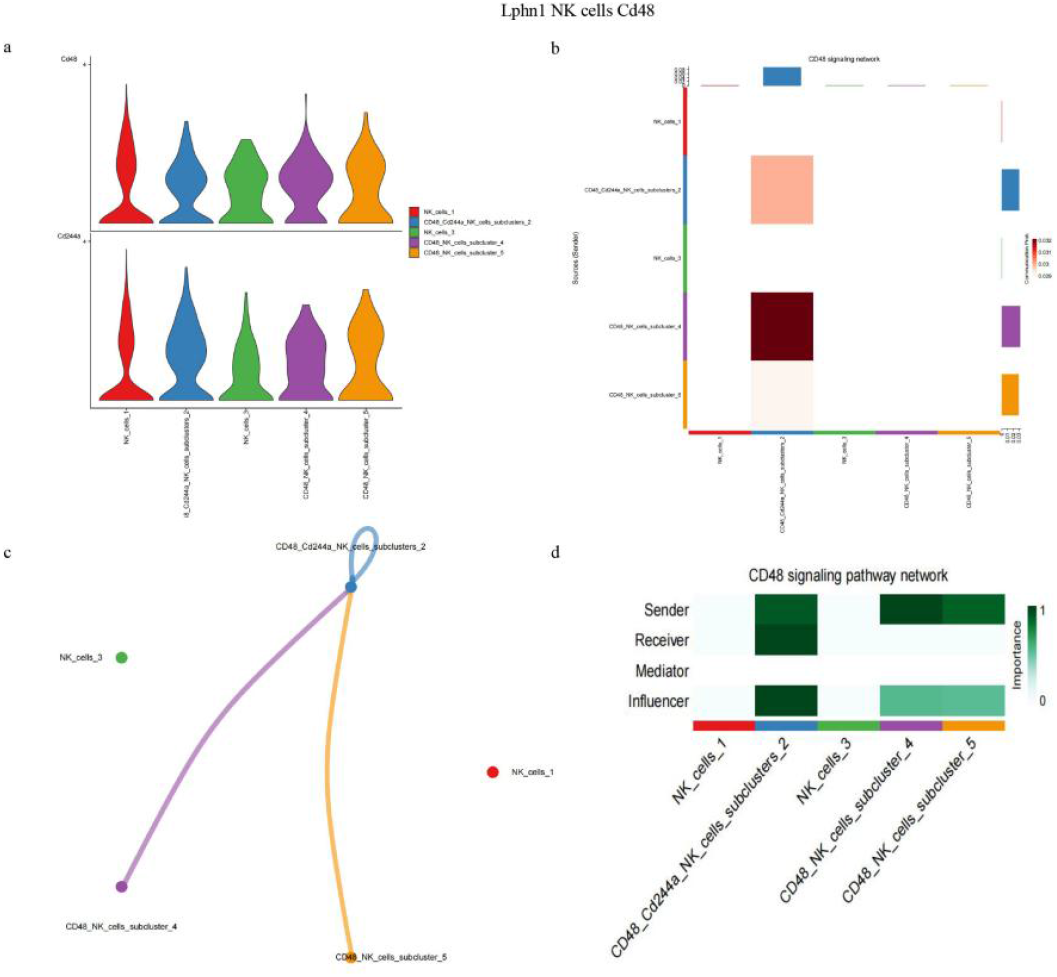
Cd48 Signaling Communication in NK Cells of the Lphn1 Group. a. Expression levels of Cd48 ligand-receptor pairs; b. Heatmap of Cd48 signaling; c. Cell communication architecture diagram of Cd48 signaling; d. Cd48 signaling communication diagram.

### 1.4 Differences in T Cells between the Lphn1 and luc Groups

Pseudotime analysis of T cells from the Lphn1 and luc groups revealed significant differences in their pseudotime cell states, as shown in Figures 9a and 9b. Notably, the cell density plot demonstrates substantial variation, indicating that there are significant functional disparities in T cells between the Lphn1 and luc groups.

**Figure 9.**
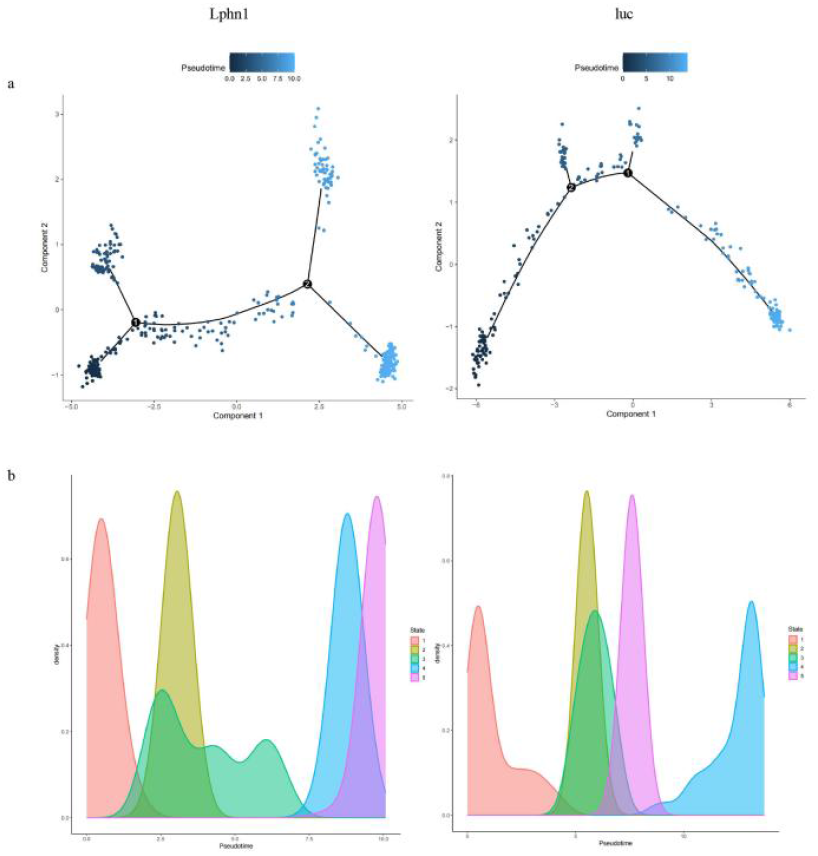
Pseudotime Analysis of T Cells in the Lphn1 and luc Groups. a. Pseudotime branching trajectory of T cells in the two groups; b. Cell state density plot of T cells in the two groups.

The cell communication analysis of T cells in the Lphn1 and luc groups revealed that Cd48 signaling was expressed only in the Lphn1 group, which is associated with anti-colorectal cancer effects, as shown in Figures 10a, 10b, 10c, and 10d. We designated the subclusters 1 and 4 of T cells in the Lphn1 group as follows: Cd244a_T_cells_subcluster_1 and Cd244a_T_cells_subcluster_4.

**Figure 10.**
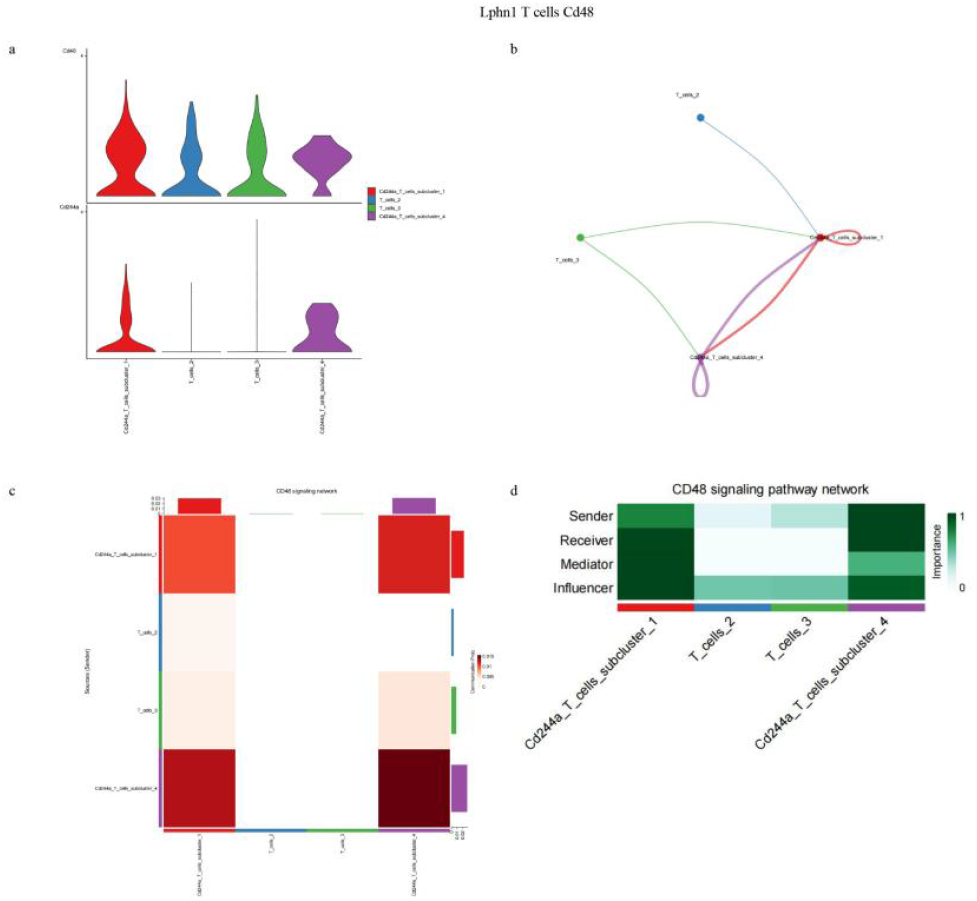
Cd48 Signaling Communication in T Cells of the Lphn1 Group. a. Expression levels of Cd48 ligand-receptor pairs; b. Cell communication architecture diagram of Cd48 signaling; c. Heatmap of Cd48 signaling; d. Cd48 signaling communication diagram.

### 2.1 Relationship Between Macrophages and T Cells

We conducted cell communication analysis between macrophages and T cells in the Lphn1 and luc groups. We found that only in the Lphn1 group was there Cd48 signaling communication between macrophages and T cells, while there was none in the luc group. Cd48 is associated with anti-colorectal cancer effects, as shown in Figures 11a, 11b, 11c, 11d, and 11e. Furthermore, as illustrated in Figures 11b, 11c, and 11e, within the cell communication between macrophages and T cells, macrophages primarily secRete Cd48 signals, while T cells receive the signals and perform downstream anti-colorectal cancer functions.

**Figure 11.**
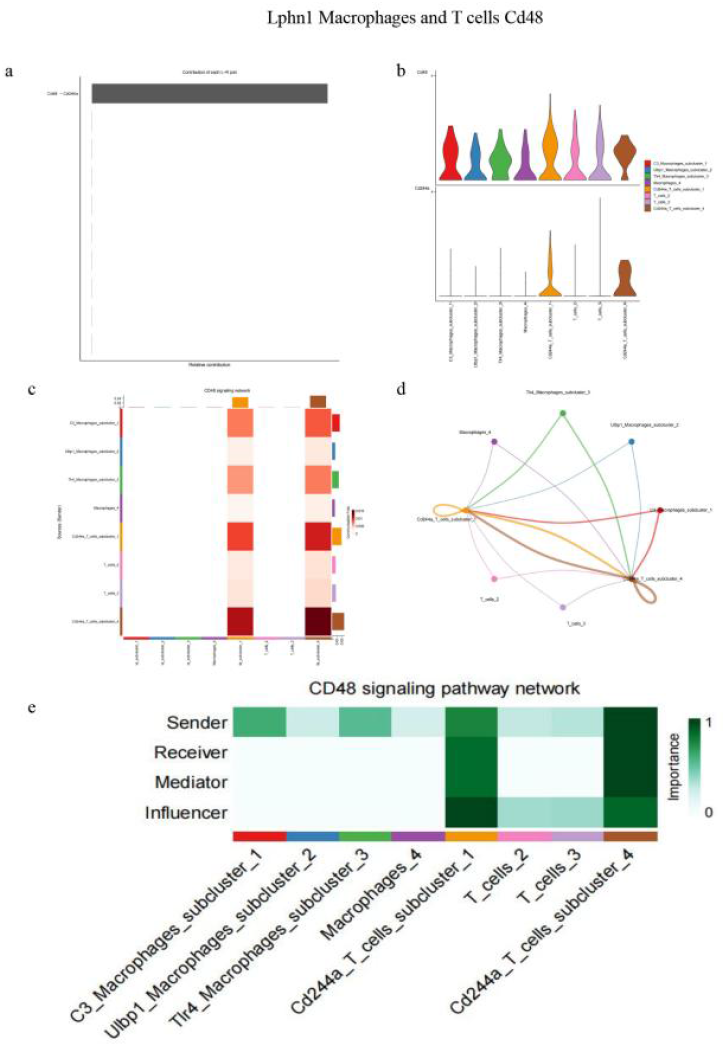
Cellular communication analysis of macrophages and T cells. a. Weight of Cd48 signaling receptor pairs; b. Gene expression levels of Cd48 receptor pairs; c. Heatmap of cellular communication analysis between macrophages and T cells; d. Structural diagram of cellular communication analysis between macrophages and T cells; e. Cellular communication analysis of combined macrophages and T cells.

### 2.2 Relationship Between CAF and Macrophages, NK Cells, and T Cells

We conducted cell communication analysis between CAF and macrophages, CAF and NK cells, as well as CAF and T cells in the Lphn1 and luc groups. The results indicate that CAFs, regardless of whether in the Lphn1 group or the luc group, possess the ability to present antigens through MHC class I signaling to activate macrophages, NK cells, and T cells, thereby enhancing their anti-tumor immune functions. This is illustrated in Figures 12a, 12b, 12c, 12d, 12e, and 12f.

**Figure 12.**
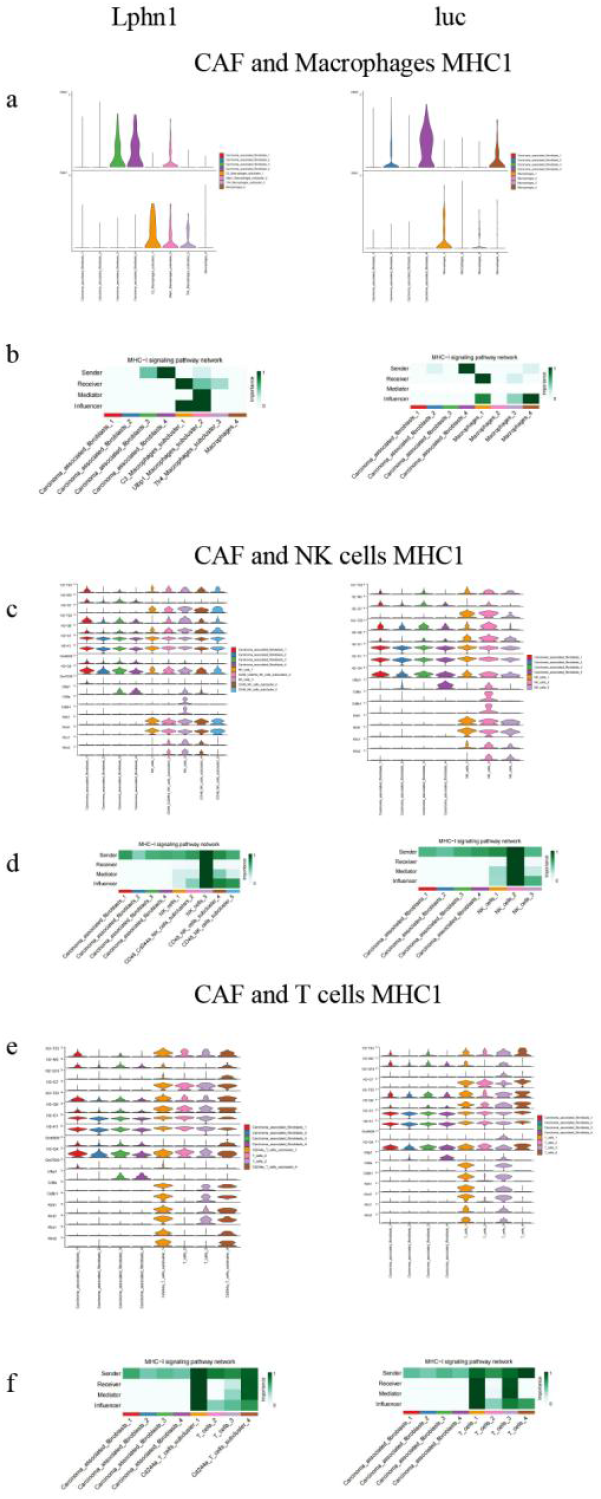
MHC Class I Signaling Pathway Cell Communication Between CAF and Macrophages, NK Cells, and T Cells. a. Gene expression levels of MHC class I signaling communication between CAF and macrophages; b. MHC class I signaling communication between CAF and macrophages; c. Gene expression levels of MHC class I signaling communication between CAF and NK cells; d. MHC class I signaling communication between CAF and NK cells; e. Gene expression levels of MHC class I signaling communication between CAF and T cells; f. MHC class I signaling communication between CAF and T cells.

### 2.3 Cell Subgroup Analysis

The KEGG enrichment results for the Cd244a_T_cells_subcluster_1 subgroup indicate that this subgroup, while receiving Cd48 signals, possesses the capability to activate the Natural Killer Cell Mediated Cytotoxicity signaling pathway to combat colorectal cancer, as shown in Figure 13a. The KEGG enrichment results for the Cd48_Cd244a_NK_cells_subcluster_2 indicate that this subgroup not only has the Cd48-Cd244a signaling but also the ability to activate the Natural Killer

**Figure 13.**
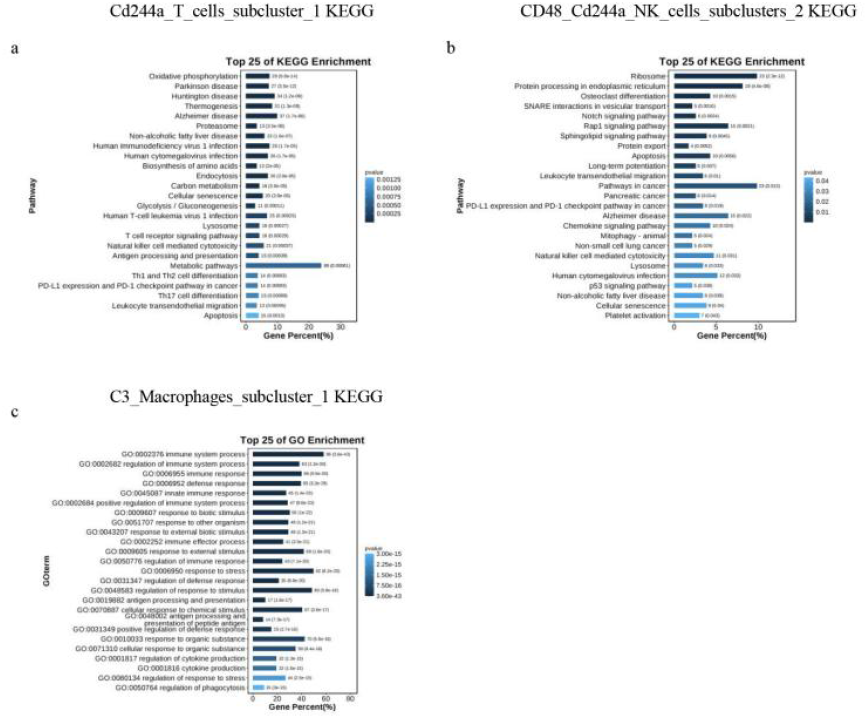
Functional Cell Subgroup GO and KEGG Enrichment Analysis. a. KEGG enrichment results for the Cd244a_T_cells_subcluster_1; b. KEGG enrichment results for the Cd48_Cd244a_NK_cells_subcluster_2; c. GO enrichment analysis for the C3_Macrophages_subcluster_1.

Cell Mediated Cytotoxicity signaling pathway against colorectal cancer, as depicted in Figure 13b. After conducting GO enrichment analysis on the C3_Macrophages_subcluster_1, we found that the functions of this subgroup include activating the following biological processes to combat colorectal cancer: positive regulation of immune system process, innate immune response, immune effector process, immune system process, regulation of immune system process, immune response, positive regulation of defense response, defense response, response to biotic stimulus, regulation of immune response, antigen processing and presentation of peptide antigen, cytokine production and regulation of phagocytosis. This is illustrated in Figure 13c.

### 2.4 High-Variance Gene Analysis

The expression level of the tumor suppressor Prdm5 in the Lphn1 knockout group is 4.5 times higher than that in the luc group, suggesting that macrophages have acquired anti-tumor immune functions after the knockout of Lphn1 (Supplementary Table 5) [19] . Vstm2a inhibits colorectal cancer and antagonizes the Wnt signaling receptor LRP6. The Vstm2a protein is significantly silenced in CRC tumor tissues and cell lines, which is mediated by high methylation of its promoter. High methylation of the Vstm2a DNA promoter and downregulation of the Vstm2a protein are associated with lower survival rates in CRC patients. Ectopic expression of Vstm2a inhibits the growth of colorectal cancer cell lines and organoids, induces apoptosis in CRC cells, and suppresses cell migration and invasion, as well as tumor growth in xenograft models in nude mice. Vstm2a is released from CRC cells via a typical secRetion pathway. The secReted Vstm2a significantly inhibits the Wnt signaling pathway in colorectal cancer cells. The Wnt signaling co-receptor low-density lipoprotein receptor-related protein 6 (LRP6) has been identified as a membrane-binding partner of Vstm2a. Through deletion/mutation and immunoprecipitation assays, we demonstrate that Vstm2a binds to the E1-4 domain of LRP6 via its IgV domain. Vstm2a inhibits LRP6 phosphorylation in a time- and dose-dependent manner, inducing LRP6 endocytosis and lysosomal-mediated degradation, collectively leading to Wnt signaling inactivation[20]. In the Lphn1 group, the expression level of Vstm2a is 3.5 times higher than that in the luc group, indicating that macrophages have gained anti-colorectal cancer functionality due to the high expression of Vstm2a after the knockout of Lphn1 (Supplementary Table 5).

In the high-variance genes of NK cells, we found that after the knockout of Lphn1, the following four genes that inhibit colorectal cancer—Ret, Oas2, Hdac11, and Ptchd4—are highly expressed: Ret (rearranged during transfection) is a transmembrane receptor tyrosine kinase and a receptor for GDNF family ligands. It acts as a tumor suppressor in colorectal cancer and shows a 5-fold increase in expression in the Lphn1 group compared to the luc group (Supplementary Table 6)[21].The invasiveness of Oas2-overexpressing RKO cells is reduced (p < 0.001–0.005). The expression level of Oas2 in the Lphn1 group is 4.4 times higher than that in the luc group.(Supplementary Table 6)[22]. Histone deacetylase (HDAC) 11 inhibits the expression of matrix metalloproteinase (MMP) 3 to suppress colorectal cancer metastasis. The expression level of Hdac11 in the Lphn1 group is 3.5 times higher than that in the luc group (Supplementary Table 6)[23]. PTCH53, a target gene of p53, is homologous to the tumor suppressor gene PTCH1 and serves as an inhibitor of Hh pathway activation. PTCH53 (formerly known as PTCHD4) exhibits strong p53 reactivity in vitro and is one of the few genes that show consistently decreased expression levels across various TP53 mutant cell lines and human tumors. Increased expression of PTCH53 can inhibit the typical Hh signaling mediated by the G protein-coupled receptor SMO. In the Lphn1 group, the expression level of PTCH53 is 3.4 times higher than that in the luc group (Supplementary Table 6)[24].

In T cells with Lphn1 knockout, three genes that exhibit anti-colorectal cancer properties can provide some mechanistic explanations for the observed suppression of colorectal cancer following Lphn1 knockout:

Chac2 is downregulated in gastric and colorectal cancers and acts as a tumor suppressor by inducing apoptosis and autophagy through the unfolded protein response. Its expression in the Lphn1 group is 5.4 times higher than that in the luc group (Supplementary Table 7)[25].Rasal2 downregulation promotes the proliferation, epithelial-mesenchymal transition, and metastasis of colorectal cancer cells. In the Lphn1 group, the expression level of Rasal2 is 5.4 times higher than that in the luc group (Supplementary Table 7)[26].Armc4/Odad2 acts as a novel negative regulator of NF-κB and a new tumor suppressor in colorectal cancer. In the Lphn1 group, the expression level of Armc4 is five times higher than that in the luc group (Supplementary Table 7)[27].

Therefore, in addition to the four target genes Ulbp1, Klrk1, Ccl6, Tlr4, Cd48, Prdm5, Vstm2a, Ret,Oas2,Hdac11 and Ptchd4. The three aforementioned subgroups should also be considered potential functional subgroups for colorectal cancer research. From a cellular perspective, the knockout of Lphn1 activates the anti-colorectal cancer functions of subpopulations of macrophages, NK cells, and T cells. Macrophages enhance antitumor immune activity by engaging the Ulbp1-Klrk1 receptor pair to activate NK cells. Additionally, macrophages activate downstream functions of T cells against colorectal cancer through CD48 signaling.

### 2.5. Transcription factor and metabolic pathway analysis

We employed pyscenic for a global transcription factor analysis of single-cell tumor-bearing mouse data stratified by cell type, as depicted in Figure 14a. Subsequently, we selected the top 9 transcription factors with the highest RSS values from each cell type, focusing on positively regulating transcription factors and those with direct interactions with the target genes identified in the aforementioned analysis. These were visually presented by cell type, as shown in Figure 14b. Figure 14c illustrates the differential expression comparison of the aforementioned transcription factors between the Lphn1 group and the luc group. Figure 14d showcases the RSS rank scoring and the display of specific genes for each cell type. Figures 14e and 14f are comprehensive expression plots and heatmaps of positively and negatively regulating transcription factors interacting with the four target genes discussed earlier, demonstrating the co-localization of the transcription factors Irf7, Nr2f1, Bclaf1, and Irf2 with the target genes Tlr4, Cd48, Prdm5, and Oas2,respectively, within specific cell types. This suggests their direct interaction and the activation of the four target genes by the transcription factors. It should be noted that Nr2f1 negatively regulates Cd48, and we observed that Nr2f1 is not expressed in the Lphn1 group, whereas Cd48 is associated with resistance to colorectal cancer. Among the four transcription factors, Irf2 inhibits colorectal cancer [32], Bclaf1 inhibits colorectal cancer [33], Irf7 inhibits colorectal cancer [34], and the function of Nr2f1 remains uncertain [35,36].

**Figure 14.**
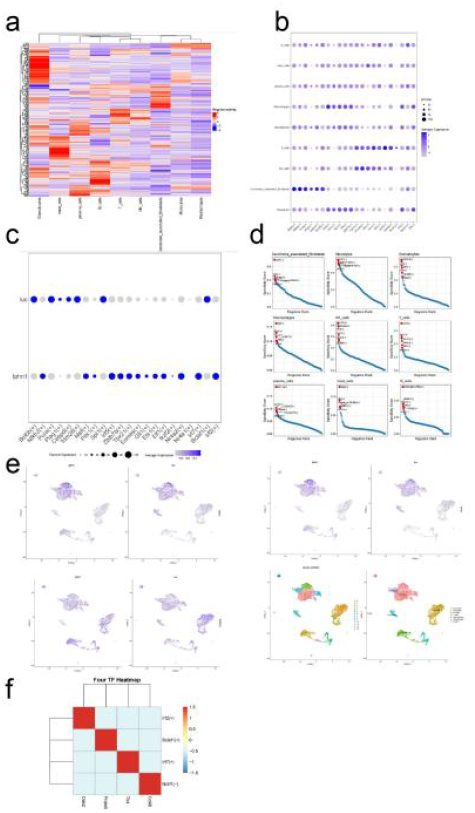
Transcription factor Analysis. a.Global transcription factor analysis of single-cell mouse data stratified by cell type; b. Top 9 transcription factors with the highest RSS values from each cell type, focusing on positively regulating transcription factors and those with direct interactions with the target genes identified in the aforementioned analysis visually presented by cell type; c. Differential expression comparison of the aforementioned transcription factors between the Lphn1 group and the luc group; d. The RSS rank scoring and the display of specific genes for each cell type; e,f. Comprehensive expression plots and heatmaps of positively and negatively regulating transcription factors interacting with the four target genes discussed earlier, demonstrating the co-localization of the transcription factors Irf7, Nr2f1, Bclaf1, and Irf2 with the target genes Tlr4, Cd48, Prdm5, and Oas2,respectively, within specific cell types.

Metabolic analysis of single-cell data from tumor-bearing mice was performed using scMetabolism. As illustrated in Figure 15a, four differentially metabolized pathways were identified between the Lphn1 group and the luc group, which are: Arginine and proline metabolism, Histidine metabolism, Phenylalanine metabolism, and Riboflavin metabolism. Among these, Arginine and proline metabolism was more active in the Lphn1 group, while Histidine metabolism was more active in the luc group. The expression of these four metabolic pathways across various cell types is shown in Figure 15b.

**Figure 15.**
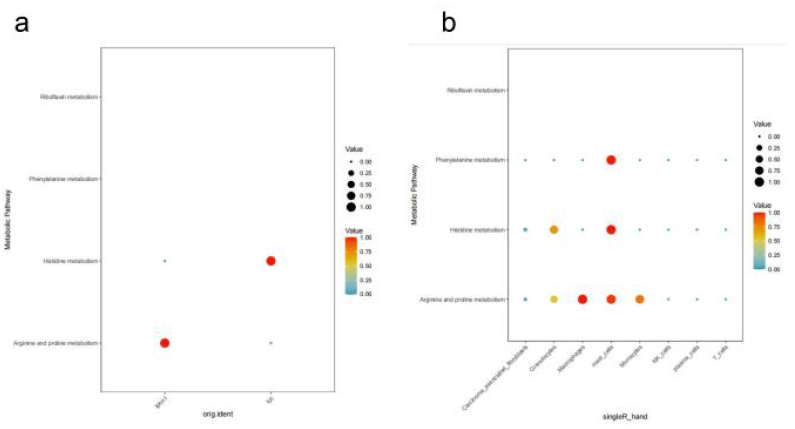
Metabolic pathway Analysis. a. Four differentially metabolized pathways were identified between the Lphn1 group and the luc group; b.The expression of these four metabolic pathways across various cell types.

Nr2f1, as a transcription factor, “indirectly negatively regulates” CD48. In previous analyses, CD48 was found to be secreted by immune cells such as NK cells, T cells, and macrophages, and its function, along with downstream signaling pathways, is to suppress colorectal cancer. This raises the suspicion that Nr2f1 may function as an oncogene and therapeutic target. Nr2f1 was significantly overexpressed in the control group, whereas its expression was markedly reduced in the tumor-suppressed group with Lphn1 knockout. Notably, Nr2f1 is predominantly expressed in cancer-associated fibroblasts (CAFs).

Nr2f1, acting as a transcription factor, “indirectly negatively regulates” CD48. Previous analyses showed that CD48 is secreted by immune cells, including NK cells, T cells, and macrophages, and that it suppresses colorectal cancer through its own and downstream signaling pathways. Thus, Nr2f1 is suspected to function as an oncogene and therapeutic target. As shown in Figure 16a, Nr2f1 was significantly overexpressed in the control group but markedly downregulated in the tumor-suppressed group with Lphn1 knockout. Additionally, Nr2f1 was predominantly expressed in cancer-associated fibroblasts (CAFs), as shown in Figure 16b.

**Figure 16.**
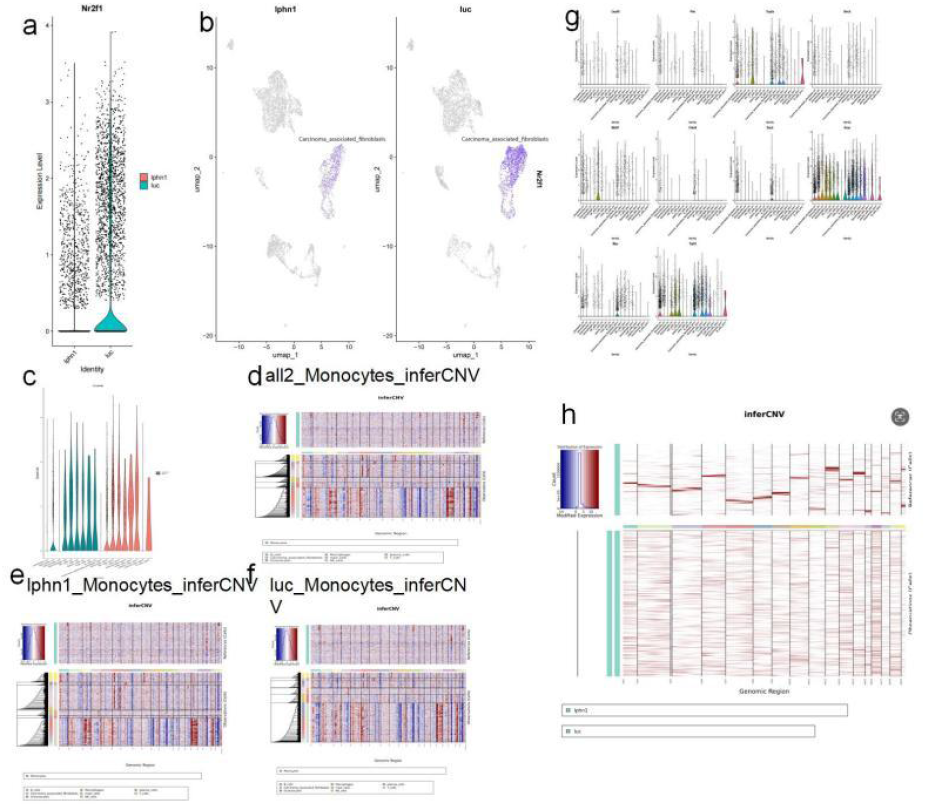
CAF originated from CT26 cells, and Nr2f1 is a potential therapeutic target. a. The expression levels of Nr2f1 in the experimental group and the control group.b. The expression status of Nr2f1 in various cell types of the experimental group and the control group.c. The expression status of CD48 in different cell types of the experimental group and the control group.d. CNVs of all cell types in the experimental group and the control group, using monocytes as a reference.e. CNVs of all cell types in the Lphn1 group, using monocytes as a reference.f. CNVs of all cell types in the luc group, using monocytes as a reference.g. The expression status of 10 CT26 marker genes in CAFs.h. Differences in CNVs of CAFs between the experimental group and the control group.

Nr2f1 “negatively regulates” the tumor-suppressive gene CD48. As shown in Figure 16c, the expression levels of CD48 did not significantly differ between the experimental and control groups. This suggests that CD48’s tumor-suppressive effects may primarily rely on downstream activated signaling pathways, consistent with previous analyses. Based on prior analyses, Nr2f1 may influence downstream signaling pathways of CD48 through certain pathways, thereby exerting antitumor functions.

CD48 showed no expression in either the experimental or control group of cancer-associated fibroblasts (CAFs), as shown in Figure 16c. This suggests that Nr2f1 may activate CD48 in the experimental group through upstream and downstream signaling pathways and paracrine signaling, thereby further activating downstream antitumor signaling pathways. I hope to clarify the specific tumor-suppressive mechanism of the Nr2f1-?-CD48-target pathway in the future through co-culture experiments, analysis, and literature review.

As shown in Figures 16d, 16e, and 16f, CAFs may be the primary proliferating malignant cells after injection of CT26 (fibroblasts). Combined with previous analyses, the number of CAFs in the weak tumor group (with Lphn1 knockout) was half that of the control group. It is therefore hypothesized that CAFs are terminally differentiated cells of CT26 and are sensitive to Lphn1 knockout. Whether this sensitivity is related to the reduction of Nr2f1 remains to be verified.

As shown in Figure 16g, the CT26 marker genes include:Top2a: DNA topoisomerase 2α, closely associated with cell proliferation.Sox2: Transcription factor Sox2, which maintains stem cell pluripotency.Myc: Proto-oncogene Myc.These three genes are highly expressed in CAFs and serve as marker genes for CT26. This provides reason to suspect that CAFs may originate from the differentiation of CT26 cells.

As shown in Figure 16h, following Lphn1 knockout, all chromosomes in CAFs exhibited overexpression.

As shown in Figures 15 and 16, CAF cells in tumors originate from CT26 cells. Additionally, Nr2f1 may serve as a potential therapeutic target, particularly as a transcription factor, for the treatment of colorectal cancer.

## Discussion

We conducted single-cell sequencing analysis on tumor-bearing mice with and without Lphn1 knockout. The analysis revealed significant differences in the quantities of various cell types. GO and KEGG enrichment analyses of differentially expressed genes indicated that macrophages restored their innate immune functions after Lphn1 knockout, while NK cells exhibited activated immune functions and enhanced anti-tumor activities. In mice with intact Lphn1, T cells in the tumors underwent programmed cell death. Pseudotime analysis showed that macrophages underwent substantial functional changes following Lphn1 knockout, acquiring the ability to positively regulate immune functions, modulate type 2 immune responses, and promote the proliferation of specific cell types, including T cells, mast cells, and epithelial cells. Cell communication analysis revealed that in macrophage subtype 2, Ulbp1 was upregulated after Lphn1 knockout, which activated the Ulbp1-Klrk1 signaling pathway, facilitating NK cell-mediated cytotoxicity (both the ligand and receptor were enriched in this pathway), thus enhancing the anti-tumor activity of NK cells. Anti-tumor macrophages were recruited by self-secReted Ccl6 (with expression levels twice that of the luc control group). Additionally, the complement system of these anti-tumor macrophages showed a reduction in a set of oncogenic ligand-receptor signals compared to the luc group. Anti-tumor macrophages secReted Tlr4 at levels 1.5 times higher than those in the luc group, leading to the transformation of macrophages into M1-type anti-tumor macrophages. Meanwhile, carcinoma-associated fibroblasts (CAFs) in tumors of Lphn1 knockout mice secReted Ccl6 at levels 0.5 times higher than those in the luc group, indicating that CAFs have the capacity to recruit anti-tumor macrophages. Pseudotime analysis revealed significant functional changes in NK cells after Lphn1 knockout. Cell communication analysis demonstrated that NK cells secReted Cd48 signals, which are associated with anti-tumor activity. Pseudotime analysis of T cells suggested that the functional differences between cells with and without Lphn1 knockout were substantial. Cell communication analysis indicated that T cells expressed Cd48 only after Lphn1 knockout, suggesting they possess anti-tumor activity. Combined cell communication analysis of macrophages and T cells showed that only after Lphn1 knockout did macrophages secRete anti-tumor Cd48 signals that were received and acted upon by T cells. Furthermore, cell communication analyses combining CAFs with macrophages, NK cells, and T cells showed that regardless of Lphn1 knockout, CAFs had the capability to present MHC1 anti-tumor signals to these three types of cells with anti-tumor potential, implying that CAFs are key antigen-presenting cells in anti-tumor activities.

Meanwhile, the high-variance gene analysis from single-cell studies identified macrophage Prdm5 and Vstm2a as potential target genes for the treatment of colorectal cancer. Therefore, we identified five target genes: Ulbp1, Klrk1, Tlr4, Ccl6,Cd48, Prdm5, Vstm2a, Ret,Oas2,Hdac11 and Ptchd4, along with three anti-tumor cell subpopulations: Cd244a_T_cells_subcluster_1, Cd48_Cd244a_NK_cells_subclusters_2, and C3_Macrophages_subcluster_1. Interventions targeting these subpopulations and the five target genes may represent new therapeutic avenues for the treatment of colorectal cancer. From a cellular perspective, the knockout of Lphn1 activates the anti-colorectal cancer functions of subpopulations of macrophages, NK cells, and T cells. Macrophages enhance antitumor immune activity by engaging the Ulbp1-Klrk1 receptor pair to activate NK cells. Additionally, macrophages activate downstream functions of T cells against colorectal cancer through CD48 signaling. After the knockout of Lphn1, macrophages are recruited by autocrine Ccl6 and Ccl6 secReted by CAFs. They exhibit high expression of Tlr4 and have the potential to transition into M1-type macrophages due to changes in their cellular state. After the knockout of Lphn1, the CAFs were reduced by half. CAFs, which are part of the cell network communication associated with tumor cells, typically play an immunosuppressive role[28-31]. A reduction by half indicates that the immunosuppression in the Lphn1 knockout group has significantly decreased. This suggests that the efficacy of various cancer immunotherapy drugs can be significantly enhanced. In specific cell types, the four colorectal cancer-resistant transcription factors Irf7, Nr2f1 (with uncertain function), Bclaf1, and Irf2 co-localize with Tlr4, Cd48, Prdm5, and Oas2, respectively, and have been analyzed by pyscenic to interact and functionally contribute to the resistance against colorectal cancer. The four differential metabolic pathways between the Lphn1 group and the luc group are Arginine and proline metabolism, Histidine metabolism, Phenylalanine metabolism, and Riboflavin metabolism, with Arginine and proline metabolism being more active in the Lphn1 group and Histidine metabolism being more active in the luc group. CAF cells in tumors originate from CT26 cells. Additionally, Nr2f1 may serve as a potential therapeutic target, particularly as a transcription factor, for the treatment of colorectal cancer.

## Conclusion

We identified five target genes for anti-colorectal cancer through single-cell analysis: Ulbp1, Klrk1, Ccl6, Tlr4, Cd48, Prdm5, Vstm2a, Ret,Oas2,Hdac11 and Ptchd4, along with their corresponding cell types. We also identified cell subpopulations with anti-tumor functions, including Cd244a_T_cells_subcluster_1, Cd48_Cd244a_NK_cells_subclusters_2, and C3_Macrophages_subcluster_1, which can be targeted for therapeutic interventions. Additionally, we determined that cancer-associated fibroblasts (CAFs) serve as the primary presenters of MHC1 antigens to macrophages, NK cells, and T cells in the fight against colorectal cancer. From a cellular perspective, the knockout of Lphn1 activates the anti-colorectal cancer functions of subpopulations of macrophages, NK cells, and T cells. Macrophages enhance antitumor immune activity by engaging the Ulbp1-Klrk1 receptor pair to activate NK cells. Additionally, macrophages activate downstream functions of T cells against colorectal cancer through CD48 signaling. After the knockout of Lphn1, macrophages are recruited by autocrine Ccl6 and Ccl6 secReted by CAFs. They exhibit high expression of Tlr4 and have the potential to transition into M1-type macrophages due to changes in their cellular state. After the knockout of Lphn1, the CAFs were reduced by half. CAFs, which are part of the cell network communication associated with tumor cells, typically play an immunosuppressive role. A reduction by half indicates that the immunosuppression in the Lphn1 knockout group has significantly decreased. This suggests that the efficacy of various cancer immunotherapy drugs can be significantly enhanced. In specific cell types, the four colorectal cancer-resistant transcription factors Irf7, Nr2f1 (with uncertain function), Bclaf1, and Irf2 co-localize with Tlr4, Cd48, Prdm5, and Oas2, respectively, and have been analyzed by pyscenic to interact and functionally contribute to the resistance against colorectal cancer. The four differential metabolic pathways between the Lphn1 group and the luc group are Arginine and proline metabolism, Histidine metabolism, Phenylalanine metabolism, and Riboflavin metabolism, with Arginine and proline metabolism being more active in the Lphn1 group and Histidine metabolism being more active in the luc group. CAF cells in tumors originate from CT26 cells. Additionally, Nr2f1 may serve as a potential therapeutic target, particularly as a transcription factor, for the treatment of colorectal cancer. These conclusions may provide new insights and avenues for the treatment of colorectal cancer, contributing to the development of personalized medicine.

## Method

### 1. Construction of Mouse Transplant Tumor Model

Cohorts of sibling mice were grouped, and 5 × 10^5^ CT26 cells (a mouse colorectal cancer cell line from ATCC) were injected into the left axilla of both BALB/c and C57BL/6J background mice. Tumor volume was measured using calipers, and the calculation formula for tumor volume was: ½ × longitudinal diameter (length) × maximum transverse diameter (width). Mice were euthanized when tumor volume reached 2000 mm^3^.

### 2. Single-Cell Sequencing Analysis

Single-cell sequencing was performed using the single-cell sequencing platform at Fudan University. Data preprocessing was carried out using Cellranger, followed by downstream analysis using Seurat. Pseudotime analysis was conducted with Monocle, and cell communication analysis was performed using CellChat. Transcription factor analysis was conducted using the software pyscenic. Metabolic analysis was performed using the software scMetabolism. CNVs were analyzed using inferCNV.

## Acknowledgement

I would like to express my gratitude to Professor Hou Xianyu and all the members of the laboratory for their assistance during the experimental process.

## Data availability

Please request the raw data from the corresponding author with a valid reason.

## Author contribution

Yi Wang design the experiment. Yi Wang conduct the experiment. Yi Wang manuscript the article.

